# Forebrain dopamine value signals arise independently from midbrain dopamine cell firing

**DOI:** 10.1101/334060

**Authors:** Ali Mohebi, Jeffrey Pettibone, Arif Hamid, Jenny-Marie Wong, Robert Kennedy, Joshua Berke

**Affiliations:** Department of Neurology, University of California, San Francisco; Department of Neuroscience, Brown University; Department of Chemistry, University of Michigan, Ann Arbor

**Author notes:** equal contributions (Ali Mohebi, Jeffrey Pettibone).

## Abstract

The mesolimbic dopamine projection from the ventral tegmental area (VTA) to nucleus accumbens (NAc) is a key pathway for reward-driven learning, and for the motivation to work for more rewards. VTA dopamine cell firing can encode reward prediction errors (RPEs^1,2^), vital learning signals in computational theories of adaptive behavior. However, NAc dopamine release more closely resembles reward expectation (value), a motivational signal that invigorates approach behaviors^3-7^. This discrepancy might be due to distinct behavioral contexts: VTA dopamine cells have been recorded under head-fixed conditions, while NAc dopamine release has been measured in actively-moving subjects. Alternatively the mismatch may reflect changes in the tonic firing of dopamine cells^8^, or a fundamental dissociation between firing and release. Here we directly compare dopamine cell firing and release in the same adaptive decision-making task. We show that dopamine release covaries with reward expectation in two specific forebrain hotspots, NAc core and ventral prelimbic cortex. Yet the firing rates of optogenetically-identified VTA dopamine cells did not correlate with reward expectation, but instead showed transient, error-like responses to unexpected cues. We conclude that critical motivation-related dopamine dynamics do not arise from VTA dopamine cell firing, and may instead reflect local influences over forebrain dopamine varicosities.

---

We trained rats in an operant, trial-and-error, “bandit” task^7^(Fig.1a,b). On each trial illumination of a nose poke port (*Light-On)* prompted approach and entry into that port (*Center-In)*. After a variable hold period (0.5-1.5s), a white noise burst (*Go Cue)* led the rat to withdraw (*Center-Out)* and poke one of the two immediately adjacent ports (*Side-In*). On rewarded trials this *Side-In* event was accompanied by an audible food hopper click, prompting the rat to collect a sugar pellet from a separate food port (*Food-Port-In).* Leftward and rightward choices were each rewarded with independent probabilities, which occasionally changed without warning. When rats were more likely to receive rewards, they were more motivated to engage in task performance. This was apparent in their “latency” – the time between *Light-On* and *Center-In -* which was sensitive to the outcome of the preceding few trials (Fig. 1c) and thereby scaled inversely with reward rate (Fig 1b).

**Figure 1:**
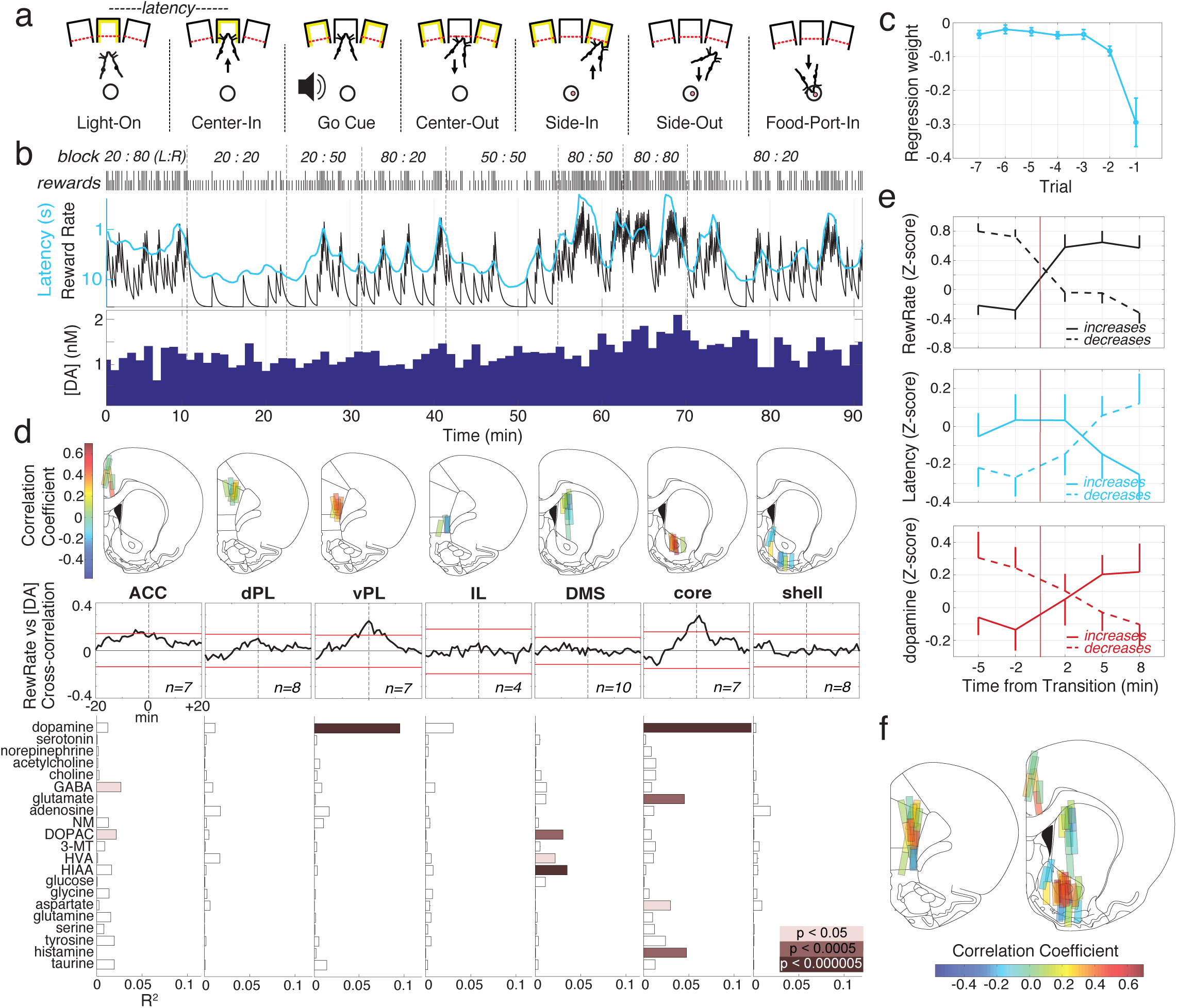
Dopamine release covaries with reward rate in NAc core and ventral prelimbic cortex. **a.** Sequence of bandit task events. **b.** Example session. Top row, reward probabilities in each block (left:right choices); Next row, tick marks indicate outcome of each trial (tall ticks, rewarded; short ticks, unrewarded). Next row, leaky-integrator estimate of reward rate (black) and running-average of latency (cyan; log scale). Bottom, NAc core dopamine time series in the same session (1 min samples). **c.** Regression analysis showing dependency of (log-) latency on the outcome of recent trials, during microdialysis sessions (n=26 sessions, 7113 trials, from 12 rats; error bars show SEM). **d.** Top, locations of microdialysis probes in medial frontal cortex and striatum. n=51 probe locations, from 12 rats, each with two microdialysis probes that were lowered further between sessions. Color of bar indicates strength of correlation between dopamine and reward rate (same data as c; see also Supplementary Fig.1). ACC, anterior cingulate cortex; dPL, dorsal prelimbic cortex; vPL, ventral prelimbic cortex; IL, infralimbic cortex; DMS, dorsal-medial striatum. Middle, average cross-correlograms between dopamine and reward rate in each region. Red bars indicate the mean 99% confidence interval generated from shuffled time series. Bottom, relationships between a range of neurochemicals and reward rate as determined through multiple regression analysis. Note that the relationship between dopamine and reward rate was highly significant in vPL and NAc core, not elsewhere (for relationships to other behavioral variables, see Supplementary Fig.2). **e.** Effect of block transitions on reward rate (top), latency (middle) and NAc core dopamine (bottom). All data is from the 14 sessions in which NAc core dopamine was measured (one per rat, combining new and previously reported^7^ data). Transitions were classified by whether the experienced reward rate increased (n=25) or decreased (n=33). Data were binned into 3 min epochs, discarding the one minute sample that included the transition time, and plotted as mean +-SEM. **f.** Composite maps of correlations between dopamine and reward rate from all microdialysis experiments (n=19 rats, 33 sessions, 58 probe placements).

We compared how dopamine firing and release vary with reward rate and motivation. First, we used microdialysis combined with liquid chromatography–mass spectrometry^9^ to simultaneously assay 21 different neurotransmitters and metabolites during bandit task performance, each with 1 min time resolution. Probes targeted seven distinct forebrain subregions within medial frontal cortex and striatum (Fig. 1d; Supplementary Fig.1). Regression analyses compared chemical time series to a range of behavioral factors (Supplementary Fig.2). We replicated our prior finding (in a different set of rats) that – unlike other neurotransmitters – mesolimbic dopamine specifically correlates with reward rate^7^ (Fig. 1d,e,f). However, we found that this relationship was localized to NAc core, and was not seen in NAc shell or dorsal-medial striatum. Similarly, dopamine release correlated with reward rate in ventral prelimbic cortex, but not in more dorsal or ventral portions of medial frontal cortex (Fig. 1d,f). This observation of twin “hotspots” of value-related dopamine release was unexpected, given that cortical and striatal dopamine are generally considered to have very different kinetics and functions^10,11^. Yet this spatiotemporal pattern has an intriguing parallel in human fMRI studies, which consistently find that BOLD signal correlates with subjective value specifically in NAc and ventral-medial prefrontal cortex^12,13^.

The NAc core receives dopamine input from lateral portions of VTA (VTA-l; ^14,15^. In head-fixed mice, VTA-l dopamine neurons reportedly have uniform, RPE-like responses to conditioned stimuli^16^. However, to our knowledge, identified dopamine cells have not been recorded in unrestrained animals performing behavioral tasks. To achieve this we used optogenetic tagging^2,17,18^ in TH:Cre rats^19^. After infecting the VTA with a virus for Cre-dependent expression of channelrhodopsin (AAV-DIO-ChR2), optrodes (Fig. 2a) were used to record single-unit responses to brief blue laser pulses (Fig 2b; Supplementary Figs. 3,4,5). Of 122 well-isolated VTA-l units, 27 showed reliable short-latency increases in firing to light onset and were considered identified dopamine neurons (see Methods). All dopamine neurons were tonically-active, with relatively low firing rates (mean 7.7Hz; range 3.7-12.9Hz; compared to the average of all VTA neurons, p<0.001 one-tailed Mann–Whitney). They also typically had longer-duration spike waveforms (compared to all VTA neurons, p<5×10^-6^, one-tailed Mann–Whitney), although there were clear exceptions (Fig. 2b), confirming prior reports that waveform duration alone is an insufficient marker of dopamine cells^2,20^. A distinct cluster of VTA neurons (n=38) had brief waveforms, higher firing rates (>20Hz; mean 41.3Hz, range 20.1-97.1Hz), and included no tagged dopamine cells. We presume that these are GABAergic and/or glutamatergic^2,21^, and refer to them as “non-dopamine” cells below.

**Figure 2:**
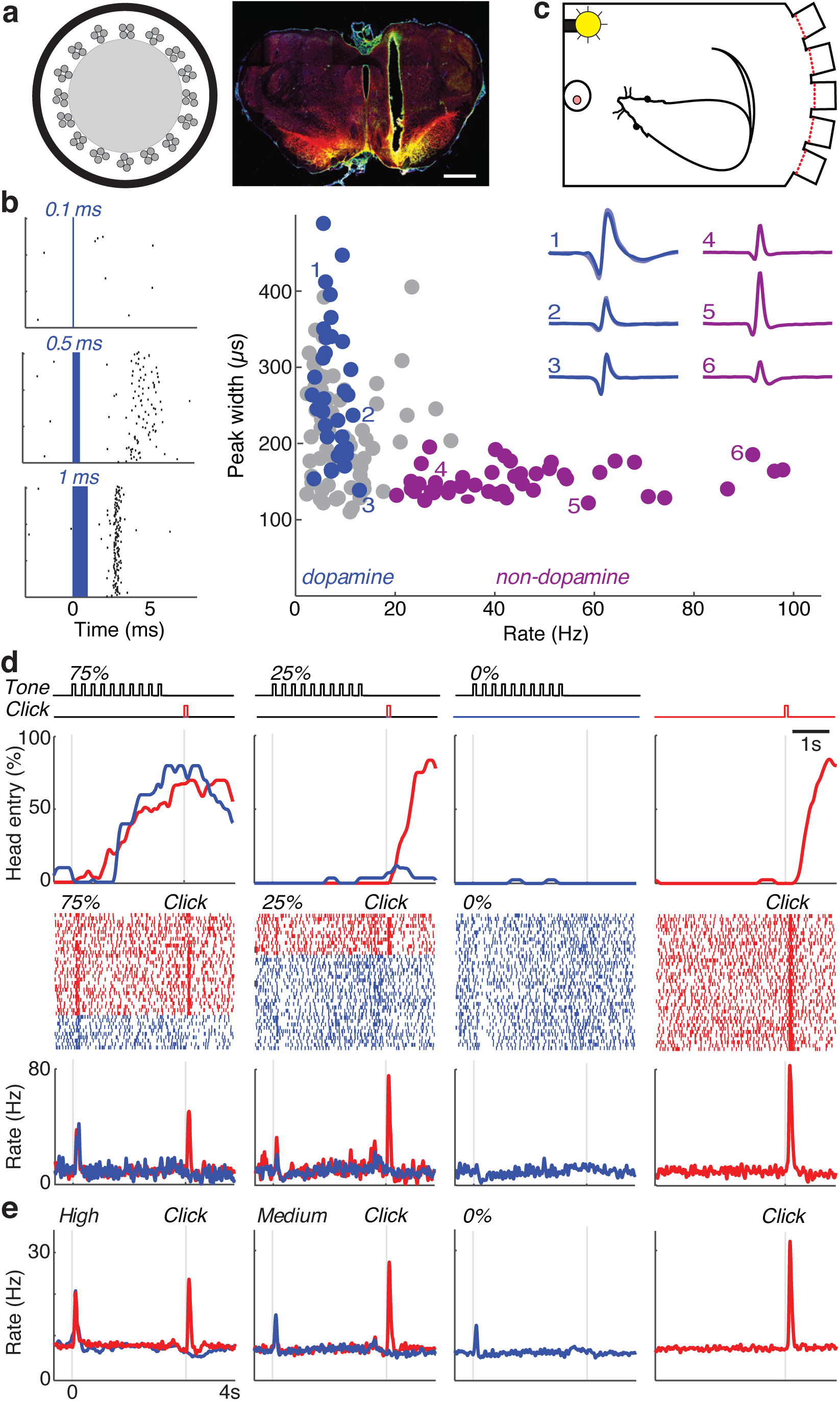
Optogenetic identification of dopamine neurons. **a.** Left, each optrode consisted of 16 tetrodes arranged around a 200µm optic fiber. Right, example of histological verification of optrode placement within lateral VTA. Scale bar = 1mm. Red = immunostaining for the dopamine cell marker tyrosine hydroxylase; green = ChR2-EYFP; yellow = overlap. For the locations of all dopamine neurons, see Supplementary Fig. 3. **b.** Left, example of optogenetic stimulation of a VTA dopamine neuron. As blue laser pulse duration increased, the neuron fired earlier and more reliably (for quantification see Supplementary Fig. 4). Right, Scatter plot of session-wide firing rate (x-axis) versus width (at half-maximum) of averaged spike waveforms for each unit. Tagged dopamine cells are in blue; purple indicates a distinct cluster of consistently untagged, presumed non-dopaminergic neurons with narrow waveforms and higher firing rate (>20Hz). Insets show examples of average waveforms (for all dopamine and non-dopamine waveforms see Supplementary Fig. 4). **c.** Pavlovian approach task was run in the same apparatus as the bandit task, but with the houselight on. **d.** Top, example of conditioned approach behavior during one Pavlovian session. “Head entry %” indicates proportion of trials for which the rat was at the food port at each moment in time. Red, blue indicate rewarded, unrewarded trials. This rat was more likely to go to the food port during the cue that was highly (75%) predictive of rewards compared to the other cues (25% and 0%); for the session shown, one-way ANOVA, F=11.1, p<1.2×10^-6^. Unpredictable reward delivery (right) prompts rapid approach. Bottom, raster plots and peri-event time histograms from an identified dopamine neuron during that same session. **e.** Averaged firing for all tagged dopamine cells (n=27) in this task. “High”/”Medium” tones were either 75%/25% predictive of reward (n=9 cells), or 100%/50% (n=18) respectively. Data on each dopamine neuron is presented in Supplementary Fig. 5.

**Figure 3:**
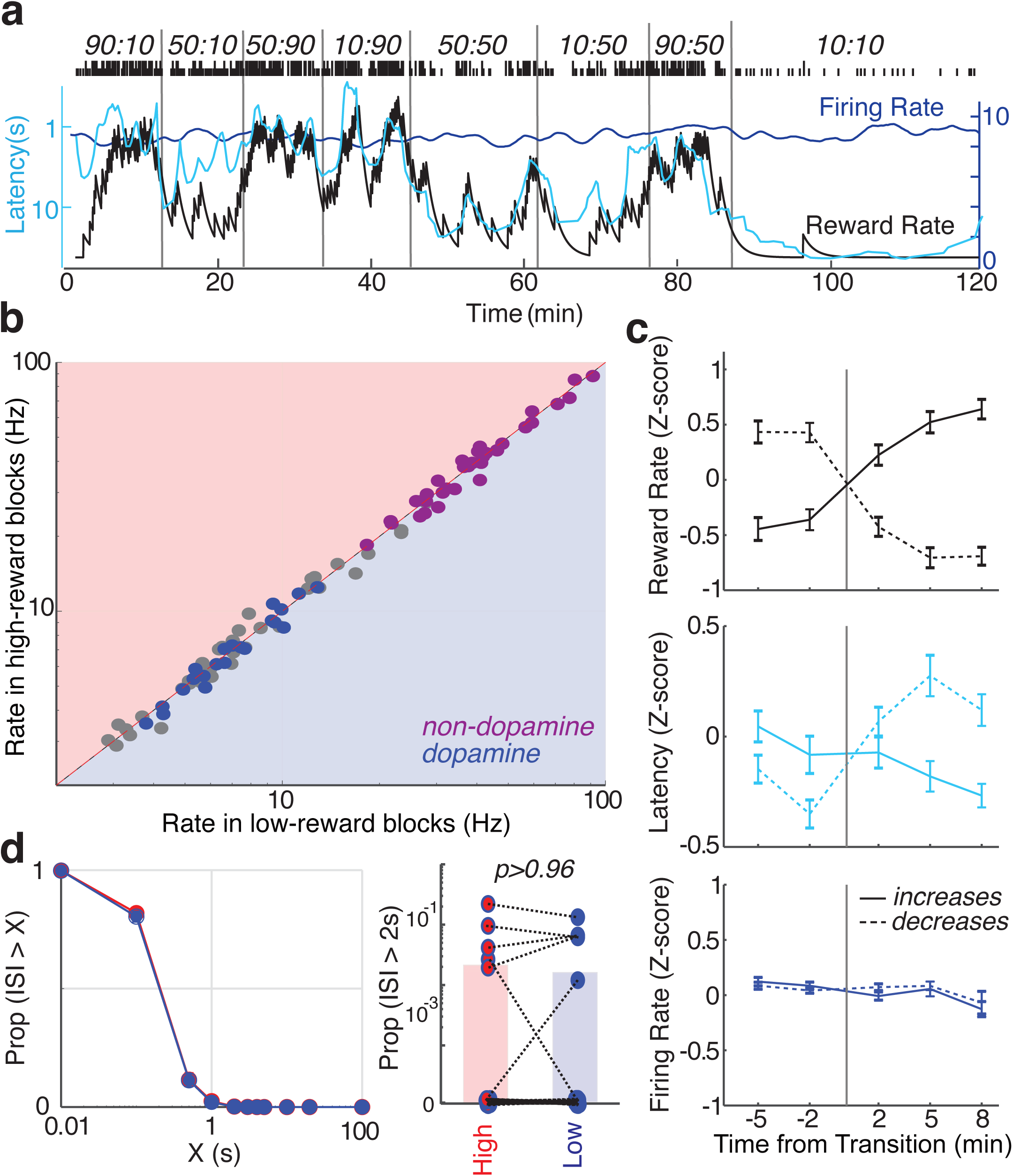
Tonic firing of dopamine cells is unrelated to motivation. **a.** Firing rate (dark blue) of one identified VTA dopamine neuron during bandit task performance. Latency (cyan) covaries with reward rate, but firing rate does not. **b**. Scatter plot showing firing rate for all VTA neurons (blue = tagged dopamine cells; purple = non-dopamine cells; grey = unclassified) in low vs high reward rate blocks. None showed significant differences in firing (Wilcoxon signed rank test using 1-min bins of firing rate, all p > 0.05 after correcting for multiple comparisons). **c.** Analysis of reward rate, latency and dopamine firing rate changes at block transitions (same format as Fig. 1e). n=95 reward rate increases and 76 decreases. **d.** Analysis of interspike intervals (ISIs). Left, overall ISI distributions are unchanged between higher- and lower reward rate blocks. Right, proportion of time spent inactive (defined as ISI > 2s) is unchanged between lower- and higher reward rate blocks. Circles indicate individual dopamine cells (n=27, same neurons as Fig.2), bars indicate mean values. Note log scale.

**Figure 4:**
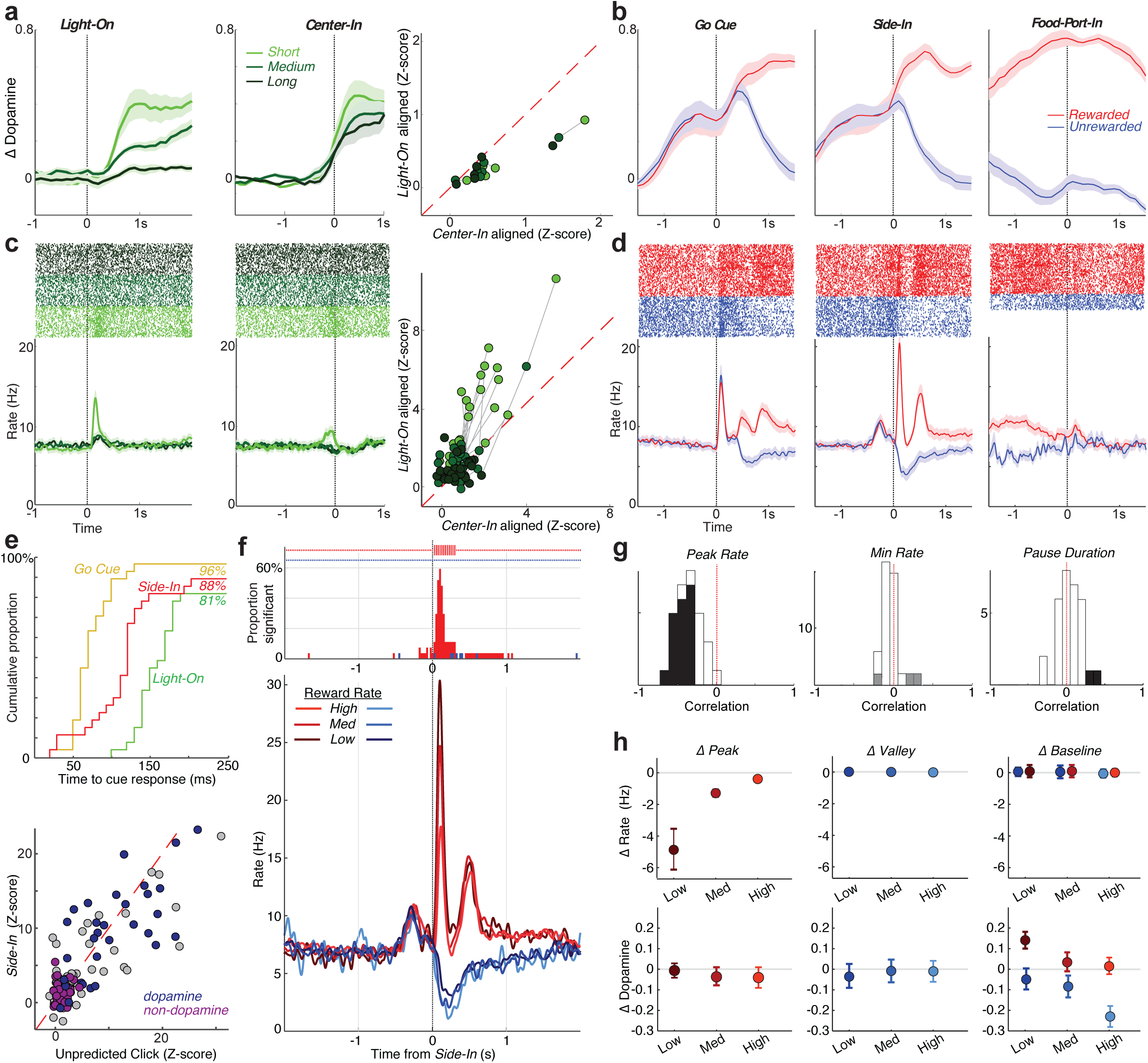
Phasic firing of VTA-l dopamine cells does not account for NAc core dopamine release. **a.** Left, event-aligned NAc core dopamine release (reanalysis of voltammetry data from ref. ^7^; n=6 rats, mean +-SEM). Green colors indicate different latencies, in equal terciles. Data are normalized by the average peak dopamine concentration for rewarded trials in each session, and shown relative to a 2s “baseline” epoch ending 1s before *Center-In*. A robust dopamine increase occurs shortly before *Center-In,* for all latencies. Right, scatter plot compares peak dopamine aligned on either *Light-On* (y-axis) or *Center-In* (x-axis). Connected lines indicate latency terciles for the same animal. Peaks were consistently larger (i.e. alignment was better) for *Center-In* (2-way ANOVA with factors of Latency and Alignment, Alignment F=3.87, p=0.05, Latency n.s. F=0.82 p=0.79). **b.** Same voltammetry data as a, aligned on later events and divided into rewarded (red) and unrewarded (blue) trials. **c,d,** As a,b but for dopamine cell spiking. Top panels show spike raster plots for one representative dopamine neuron, bottom panels show firing rate averaged over all dopamine cells. Connected lines in the scatter plot indicate latency terciles for the same neuron. A *Light-On* response is preferentially seen for short-latency trials; this same cue-evoked response appears smaller and spread-out in the *Center-In* alignment (2-way ANOVA with factors of Latency and Alignment, Alignment x Latency interaction F= 7.47, p=0.0008). No other dopamine cell firing increase is visible at *Center-In*. **e.** Top, Cumulative distributions of time taken for dopamine cells to significantly increase firing following each of three cue onsets (*Light-On, Go-Cue,* rewarded *Side-In*). For *Light-On*, only the short-latency tercile was included. The slower response to *Light-On* is consistent with prior reports^55^ that visual cues take longer to evoke dopamine cell firing compared to auditory cues. Bottom, scatter plot comparing cell firing responses to the food hopper click in the Pavlovian task (x-axis, unpredicted trials only) and the bandit task (y-axis). As in Fig.2, dopamine cells are in blue, non-dopamine in purple, unclassified in grey. Non-dopamine cells were generally indifferent to cue onsets, instead increasing firing in conjunction with movements (Supplementary Fig. 7). **f.** Reward expectation affects dopamine cell firing during, but not before, reward feedback. Lower panel shows average dopamine cell firing rate relative to *Side-In* event, broken down by reward rate (terciles, calculated separately for each neuron then averaged). Upper plots show the fraction of individual dopamine cells whose firing rate significantly varies with reward rate at each moment in time, with bold marks indicating a proportion significantly higher than chance (binomial test, p < 0.01). Data are separated into rewarded (red) and unrewarded (blue) trials. Note that before *Side-In* dopamine cell firing did not depend on reward rate. At *Side-In,* the dopamine cell response to the reward click was much stronger when reward rate was low, consistent with RPE coding (lower reward expectation = larger prediction error if the reward cue arrives). When the reward click was omitted dopamine cells transiently reduced firing. **g.** Correlations between reward rate and individual dopamine cell peak firing rate (within 250ms after rewarded *Side-In*), minimum firing rate (middle; within 2s after unrewarded *Side-In*), and pause duration (bottom; maximum inter-spike-interval within 2s after unrewarded *Side-In*). For all histograms, grey indicates cells with significant correlations (p < 0.01) before multiple comparisons correction, black indicates cells that remained significant after correction. Positive RPE coding is strong and consistent, but negative RPE coding is weak. **h.** Comparison between consecutive trials shows that an unexpected rewarded trial (one that occurs when reward rate has been low) causes peak dopamine cell firing to be substantially diminished on the next trial, but has no effect on firing rate earlier in the trial (“baseline”, -3s to -1s relative to *Center-In).* This is consistent with positive RPE coding. By contrast, unexpected rewards do not reduce peak NAc core dopamine release on the next trial. Instead they increase “baseline”, consistent with value coding.

Recordings were typically stable for many hours, allowing us to examine activity patterns of the same individual dopamine cells across multiple behavioral tasks. For better comparison with previous work we first show responses to unpredicted food delivery and Pavlovian conditioned cues. A series of tone pips were followed by reward delivery with different probabilities (zero, medium, high) depending on the tone pitch. During prior training rats had learned about these different probabilities, as indicated by their corresponding scaled likelihood of entering the food port during cue presentation (Fig. 2d). The three different cues, together with occasional unheralded food deliveries, were given in random, interleaved order (with inter-trial interval of 15-30s). Identified dopamine neurons responded most strongly to unanticipated food hopper clicks, and progressively less strongly when these clicks were preceded by the medium-probability and high-probability cues (Fig. 2d,e). Conversely, at cue onset dopamine cells responded most strongly to the high-probability cue, and progressively less strongly to the medium- and zero-probability cues. This pattern is broadly consistent with prior reports^22-24^, confirming that VTA-l dopamine cells can display canonical RPE-like coding even in unrestrained animals.

We then turned to the bandit task. Based on evidence from anesthetized animals, it has been argued that altered dopamine levels measured with microdialysis arise from changes in the tonic firing rate of dopamine cells, and/or the proportion of active versus inactive dopamine neurons^8,25^. We therefore assessed whether these factors vary with reward rate, in a manner that could account for our microdialysis observations.

Unlike forebrain dopamine release, tonic dopamine cell firing in each block of trials was strikingly indifferent to reward rate (Fig. 3a). There was no significant change in the firing rates of individual dopamine cells – or any other VTA-l neurons - between higher- and lower-reward blocks (Fig. 3b,c; see also ref. ^26^ for concordant results in head-fixed mice). Furthermore, we never observed any dopamine cells switching between active and inactive states. The proportion of time that dopamine cells spent in long inter-spike-intervals was very low, and did not change between higher- and lower-reward blocks (Fig. 3d). Nor was there any overall change in the rate at which dopamine cells fire bursts of spikes (Supplementary Fig. 6). We conclude that changes in tonic VTA dopamine cell firing are not responsible for the motivation-linked changes in forebrain dopamine release observed in this task.

We next considered fluctuations in dopamine cell firing around specific bandit task events. Several groups have found that motivated approach behaviors are accompanied by rapid increases in NAc core dopamine, on a sub-second to seconds timescale^3-6^. In this specific task we find^7^ that NAc core dopamine rapidly increases as rats initially approach *Center-In* (Fig, 4a), and increases further towards the end of rewarded trials as they approach *Food-Port-In* (Fig. 4b). The initial increase is better aligned on *Center-In* than *Light-On* (in 6/6 voltammetry animals; for individual animal data see ref ^7^), and occurs for all latencies (Fig. 4a).

The pattern of VTA-l dopamine cell firing was very different (Fig.4c,d). The *Light-On, Go-Cue,* and (on rewarded trials) *Side-In* events all produced fast increases in the activity of most dopamine neurons (Fig. 4e). Critically, these firing changes were best aligned to the sensory cues, rather than the behaviors they evoked. Twenty-two VTA dopamine neurons significantly increased firing after *Light-On*; in all cases this response was better aligned to *Light-On* than *Center-In,* and was largest for short-latency trials (Fig 4c). No separate increase in dopamine cell firing was apparent around *Center-In,* either at the population level (Fig. 4c) or individually (Supplementary Fig.5). During the subsequent approach to *Food-Port-In* dopamine cell firing was more variable (Supplementary Fig. 5) but again showed no overall increase (Fig. 4d).

The response of dopamine cells to the reward cue at *Side-In* depended on recent reward history, in a manner consistent with RPE coding. When reward rate was low (i.e. rats had lower expectation of reward), dopamine cells responded strongly, but this response was greatly blunted when reward rate was high (Fig. 4f,g). These RPE-like responses appeared very similar if reward expectation was estimated in other ways, including trial-based reinforcement learning models (actor-critic or Q-learning) or simply counting the number of rewards in the last 10 trials (Supplementary Fig. 8). RPE coding on unrewarded trials was also visible, but much less robust (Fig. 4f). None of the 27 dopamine neurons showed a significant individual correlation between reward rate and the minimum firing rate following reward omission (Fig. 4g; all p > 0.01 after multiple comparisons correction). It has been proposed that negative RPEs may be encoded in the duration of dopamine cell pauses^27^, but this was observed in just 2/27 individual neurons (Fig. 4g, right). Such asymmetric RPE coding has been observed before^22,28,29^ and provides another dissimilarity to NAc dopamine release, which shows a larger and more sustained decrease after more disappointing outcomes^7,30^.

Despite the influence of reward expectation over the rats’ motivation to perform the task (Fig. 3c) dopamine cell firing was not dependent on reward expectation until rats heard the reward cue at *Side-In* (Fig. 4f; Supplementary Fig. 8). To further compare the impact of reward history on dopamine firing versus release we employed a consecutive-trial analysis^7^. Receiving a reward had no impact on “baseline” dopamine cell firing rates early in the subsequent trial (Fig. 4h). Instead, it reduced the magnitude of the peak response to a subsequent reward cue, consistent with (positive) RPE coding. This pattern is unlike NAc core dopamine release. The peak NAc core dopamine response to the reward cue is unchanged by reward on the preceding trial (Fig. 4h), consistent with encoding value rather than RPE^7^. Overall, we conclude that VTA-l dopamine cell firing does not account for the motivation-related patterns of dopamine release we observe in NAc core.

Spiking of VTA dopamine cells is undoubtedly important for NAc dopamine release^31^. However, local receptors on NAc dopamine terminals also powerfully modulate release^32-35^, even when VTA spiking is suppressed^36,37^. It has been noted for decades that these two dopamine control mechanisms might serve different functional roles in behavior^32,38^. However, to our knowledge midbrain dopamine cell firing and forebrain dopamine release have not previously been compared in the same behavioral situation. Our results here demonstrate that recording dopamine cells is not sufficient for understanding the functional information conveyed by dopamine transmission.

VTA-l provides the predominant source of dopamine to NAc core^14,15^. As shown here and elsewhere^16^ VTA-l dopamine cells have relatively uniform, RPE-like activity patterns, but there is increasing evidence that other dopamine subpopulations may carry distinct signals^18,39,40^. We cannot rule out the possibility that firing of dopamine cell subpopulations not recorded from here is responsible for value-related dopamine release in NAc core. However, value-related firing has never been reported for any dopamine cells, across a wide range of studies in rodents^26^ and non-human primates. It also seems unlikely that an unidentified dopamine subpopulation would dominate NAc core release, overwhelming the extensive, well-characterized input from VTA-l.

Our findings argue against the idea^25^ that increased “tonic” dopamine cell firing is responsible for increased mesolimbic dopamine with motivational arousal, as measured with microdialysis^41^. Although tonic firing can be altered by lesions or drug manipulations^8^, we are not aware of evidence that dopamine cell firing shows prolonged changes under any natural behavioral condition. Firing has been seen to ramp downwards on a ∼1s timescale as monkeys anticipate motivationally-relevant events^42,43^. However, this decline is the opposite of what would be required to boost dopamine release with reward expectation, and instead seems more akin to a sequence of transient negative prediction errors^44^.

It has been suggested that dopamine release arising from RPE-coding phasic dopamine bursts could temporally summate^45^, resulting in a tonic dopamine signal that encodes reward rate (just like the leaky integrator metric we used here). There are several reasons to think this is not the case. First, increases in NAc core dopamine after unpredicted rewards are highly transient (subsecond duration^46^), consistent with efficient clearing of dopamine from the extracellular space^7,47^. Second, we observed no overall difference in the rate of dopamine cell bursting between higher- and lower-reward blocks, suggesting that bursts are not an effective way of tracking rewards over time. Finally, although dopamine release may be particularly driven by bursts in anesthetized animals^47^, to our surprise during the bandit task we observed the opposite. Burst firing of VTA dopamine cells does produce transient NAc core dopamine increases, but these are de-emphasized in favor of the ramps that accompany approach behaviors.

How closely dopamine release within a particular forebrain area corresponds to midbrain dopamine cell firing likely depends on the specific behavioral context. Distinct striatal subregions contribute to different types of decisions, and may influence their own dopamine release according to need^48^. The NAc core is not needed for highly-trained behavioral responses to conditioned stimuli^49-51^ but is particularly important when deciding to perform time-consuming work to obtain rewards^52^. NAc core dopamine appears to provide an essential dynamic signal of how worthwhile it is to allocate time and effort to work^7,48^, even though this signal is not present in dopamine cell firing.

## Acknowledgements

We thank Peter Dayan, Loren Frank, Chris Donaghue, and Thomas Faust for their comments on an early version of the manuscript, and Rahim Hashim for technical assistance. This work was supported by the National Institute on Drug Abuse, the National Institute of Mental Health, the National Institute on Neurological Disorders and Stroke, the University of Michigan Ann Arbor, and the University of California San Francisco.

## Contributions

A.M. performed and analyzed the electrophysiology, J.P. performed and analyzed the microdialysis, and A.H. performed and analyzed the voltammetry. Microdialysis procedures were assisted by J.W. and supervised by R.K. J.D.B. designed and supervised the study, and wrote the manuscript.

## Methods

### Animals

All animal procedures were approved by the University of Michigan or University of California San Francisco Institutional Committees on Use and Care of Animals. Male rats (300– 500g, either wild-type Long-Evans or TH-Cre+ with a Long-Evans background^19^) were maintained on a reverse 12:12 light:dark cycle and tested during the dark phase. Rats were mildly food deprived, receiving 15 g of standard laboratory rat chow daily in addition to food rewards earned during task performance.

### Behavior

Pretraining and testing were performed in computer-controlled Med Associates operant chambers (25 cm × 30 cm at widest point) each with a five-hole nose-poke wall, as previously described^7^. Bandit task sessions used the following parameters: block lengths were 35-45 trials, randomly selected for each block; hold period before Go cue was 500-1500 ms (uniform distribution); left/right reward probabilities were 10,50,90% (electrophysiology rats, and previously reported^7^ voltammetry and microdialysis rats), or 20,50,80% (newly reported microdialysis rats). Electrophysiology rats also performed a Pavlovian approach task immediately after the bandit task, in the same operant chamber with the houselight on throughout the session. Three auditory cues (2 kHz, 5 kHz, 9 kHz) were associated with different probabilities of food delivery (counterbalanced across rats). Cues were played as a train of tone pips (100 ms on / 50 ms off) for a total duration of 2.6 s followed by a delay period of 500ms. Cues, and unpredicted reward deliveries, were delivered in pseudorandom order with a variable inter-trial interval (15-30 s, uniform distribution).

Current reward rate was estimated using a simple, time-based leaky-integrator^53^. Reward rate was incremented each time a reward was received, and decayed exponentially at a rate set by parameter τ (the time in s for the reward rate to decrease by ∼63%, 1-1/e). For all analyses, τ was selected based on the rat’s behavior, maximizing the (negative) correlation between reward rate and log(latency) in each session. The correlations between forebrain dopamine and reward rate were not highly sensitive to this choice of τ (Supplementary Fig. 1).

To classify block transitions as “increasing” or “decreasing” in reward rate, we compared the average leaky-integrator reward rate in the last 5 min of a block to the average reward rate in the first 8 min of the subsequent block.

### Microdialysis

#### Surgery

Rats were implanted bilaterally with guide cannula (CMA, #830 9024) in cortex and striatum. One group (n=8) received one guide cannula targeting prelimbic and infralimbic cortex (AP +3.2 mm, ML 0.6 mm relative to bregma; DV 1.4 mm below brain surface) and another targeting dorsomedial striatum and nucleus accumbens in the opposite hemisphere (AP +1.3, ML 1.9, DV 3.4). Both implants were angled 5 degrees away from each other along the rostral-caudal plane. A second group (n=4) received one guide cannula targeting anterior cingulate cortex (AP +1.6, ML 0.8, DV 0.8) and another targeting accumbens (core/shell in the opposite hemisphere at AP +1.6, ML 1.4, DV 5.5 (n=2) or AP +1.6, ML 1.9, DV 5.7 (n=2). Implant sides were counterbalanced across rats. Animals were allowed to recover for 1 week prior to retraining.

#### Chemicals

Water, methanol, and acetonitrile for mobile phases were Burdick & Jackson HPLC grade, purchased from VWR (Radnor, PA). All other chemicals were purchased from Sigma Aldrich (St. Louis, MO) unless otherwise noted. Artificial cerebrospinal fluid (aCSF) was comprised of 145 mM NaCl, 2.68 mM KCl, 1.40 mM CaCl_2_, 1.01 mM MgSO_4_, 1.55 mM Na_2_HPO_4_, and 0.45 mM NaH_2_PO_4_, adjusted pH to 7.4 with NaOH. Ascorbic acid (250 nM final concentration) was added to reduce oxidation of analytes.

#### Sample Collection and HPLC-MS

On testing day, animals were placed in the operant chamber with the houselight on. Custom-made concentric polyacrylonitrile membrane microdialysis probes (1 mm dialyzing AN69 membrane; Hospal, Bologna, Italy) were inserted bilaterally into guide cannula and perfused continuously (Chemyx Inc., Fusion 400) with aCSF at 2 µL/min for 90 min to allow equilibration. After 5 min baseline collection the houselight was extinguished, cueing the animal to bandit task availability. Sample collection continued at 1 min intervals and samples were immediately derivatized with 1.5 µL sodium carbonate, 100 mM; 1.5 µL BzCl, 2% (v/v) BzCl in acetonitrile; and 1.5 µL isotopically labeled internal standard mixture diluted in 50% (v/v) acetonitrile containing 1% (v/v) sulfuric acid, and spiked with deuterated ACh and choline (C/D/N isotopes, Pointe-Claire, Canada) to a final concentration of 20 nM. Sample series collection alternated between the two probes at 30-second intervals in each of 26 sessions, except for one session in which a broken membrane resulted in just one series (51 sample series total). Samples were analyzed using Thermo Fisher Accela UHPLC system or Thermo Fisher Vanquish UHPLC interfaced to a Thermo Fisher TSQ Quantum Ultra triple quadrupole mass spectrometer fitted with a HESI II ESI probe, operating in multiple reaction monitoring. Five µL samples were injected onto a Phenomenex core-shell biphenyl Kinetex HPLC column (2.1 mm x 100 mm). Mobile phase A was 10 mM ammonium formate with 0.15% formic acid, and mobile phase B was acetonitrile. The mobile phase was delivered an elution gradient at 450 µL/min as follows: initial, 0% B; 0.01 min, 19% B; 1 min, 26% B; 1.5 min, 75% B; 2.5 min, 100% B; 3 min, 100% B; 3.1 min, 5% B; and 3.5 min, 5% B. Thermo Xcalibur QuanBrowser (Thermo Fisher Scientific) was used to automatically process and integrate peaks. Each of the >100,000 peaks were visually inspected to ensure proper integration.

#### Analysis

All neurochemical concentration data were smoothed with a 3-point moving average (y’ = [0.25*(y-1) + 0.5(y) + 0.25*(y+1)]) and z-score normalized within each session to facilitate between-session comparisons. For each target region, a cross-correlogram was generated for each session and the average of the sessions was plotted. 1% confidence boundaries were generated for each subplot by shuffling one time series 100,000 times and generating a distribution of correlation coefficients for each session. Multiple regression models were generated using the *regress* function in MATLAB, with the neurochemical as the outcome variable and behavioral metrics as predictors. Regression coefficients were determined significant at three alpha levels (0.05, 0.0005, 0.000005), after Bonferroni-correction for multiple comparisons (alpha0020/ (21 chemicals * 7 regions * 9 behavioral regressors)).

### Electrophysiology

Rats (n=23) were implanted with custom designed drivable optrodes, each consisting of 16 tetrodes (constructed from 12.5µm nichrome wire, Sandvik, Palm Coast, FL) glued onto the side of a 200µm optic fiber and extending up to 500µm below the fiber tip. During the same surgery, we injected 1µl of AAV2/5-EF1a-DIO-ChR2(H134R)-EYFP into the lateral VTA (AP - 5.6, ML 0.8, DV 7.5). Wideband (1-9000Hz) brain signals were sampled (30,000 samples/s) using Intan digital headstages. Optrodes were lowered at least 80µm at the end of each recording session. Individual units were isolated offline using a MATLAB implementation of MountainSort^54^ followed by careful manual inspection.

#### Classification

To identify whether an isolated VTA-l unit was dopaminergic (TH+), we used the stimulus-associated latency test^17^. Briefly, at the end of each experimental session, we connected the optrode through a patch cable to a laser diode and delivered light pulse trains of different widths and frequencies. For a unit to be identified as light responsive it needed to reach the significance level of p<0.001 for 5ms and 10ms pulse trains. We also compared the light evoked waveforms (within 10ms of laser pulse onset) to session-wide averages; all light-evoked units had a Pearson correlation coefficient of >0.9. Dopamine neurons were successfully recorded from four rats (IM657, 1 unit; IM1002, 3 units; IM1003, 15 units; IM1037, 9 units). Peak width was defined as the full-width-at-half-maximum of the most prominent negative component of the aligned, averaged spike waveform. Non-tagged VTA neurons with session-wide firing rate > 20 Hz and peak width < 200 µs were classified as non-dopamine cells. To ensure that we were comparing dopamine and non-dopamine cells within the same subregions, we only analyzed non-dopamine cells recorded during sessions with at least one optically-tagged dopamine cell.

#### Analysis

For comparison of “tonic” firing to reward rate, dopamine spikes were counted in 1 min bins. To examine faster changes, spike density functions were constructed by convolving spike trains with a Gaussian kernel with variance 20 ms. To determine how quickly a neuron responded to a given cue, we used 40 ms bins (sliding in steps of 20 ms) and used a shuffle test (10,000 shuffles) for each time bin comparing the firing rate after cue onset to firing rate in the 250 ms immediately preceding the cue. The first bin at which the post cue firing rate was significantly (p<0.01, correcting for multiple comparisons) greater than baseline firing was considered the time to cue response. Peak firing rate was calculated as the maximum (Gaussian-smoothed) firing rate of each trial in a 250 ms window after *Side-In* for rewarded trials, and the valley was calculated as the minimum firing rate in a 2 s window, starting one second after *Side-In* for unrewarded trials. To compare firing rates in “high” and “low” reward blocks, for each session we performed a median split of average leaky-integrator reward rate in each block.

### Voltammetry

Fast-scan cyclic voltammetry results shown here reanalyze data previously presented and described in detail^7^.

### Data and Code Availability

All data is available *[at time of publication]* through the Collaborative Research in Computational Neuroscience (CRCNS.org) data sharing website. Custom MATLAB code is available upon request to J.D.B.

## Figure Legends

**Supplementary Figure 1. a.** Anatomical definitions of the subregions examined with microdialysis are shown at top left. Atlas sections are from ^56^. The remaining sections map the correlation between dopamine release and reward rate at individual probe placements in coronal (mm from bregma, B) and sagittal (mm from midline) planes. Color bar shows strength of correlation., **b.** Dependence of the correlation between dopamine and reward rate on the time constant (tau) of the leaky integrator used to define reward rate. As illustrated in the top panel, a larger tau indicates integration over longer time periods. Below, the dopamine: reward rate correlation evolves as a function of tau. In main figures tau was chosen (from a range of 1-1200s) to maximize the (negative) correlation between reward rate and (log) latency in each session. Thin lines represent individual sessions, with the best fit tau used in regression analyses indicated by a dot. Thick lines indicate the average of all dopamine: reward rate correlations for a given tau within each subregion.

**Supplementary Figure 2. Correlations between all neurochemicals and a range of behavioral factors.** Bars represent R^2^ values for linear tests between each analyte (rows) and behavioral covariates (columns). In models with more than one covariate, bar length indicates the R^2^ for the full model. Negative relationships are reported in blue and positive relationships are in red. P-values are reported at three alpha levels (0.05, 0.0005, 0.000005) and were Bonferroni corrected for multiple comparisons (7 subregions x 21 analytes x 12 measures). To calculate reward rate, we averaged the leaky-integrator-estimated reward rate in 1 min bins defined by the start and end of each dialysis sample. ‘Attempts’ is the number of initiated trials (including trials that resulted in an error) in each dialysis minute. Attempts and reward rate and an interaction term were combined in a single model (column 2) to examine whether adding attempts could explain additional variance in the analyte signal that could not be explained by reward rate alone. “Latency” is the average of the (log)-latency in each minute. ‘Exploit’ is the proportion of choices of the higher reward probability option, in the last half of blocks for which the two ports had different probabilities. ‘Rewards’ and ‘Omissions’ were defined as the number of rewarded and unrewarded trials in each min, respectively. ‘Cumulative Rewards’ and ‘Time’ were included in the same regression model to estimate progressive factors such as satiety, and possible slow timescale increases or decreases in analyte concentration across the session. Cumulative Rewards represents the total number of rewards received by the end of the current dialysis minute, and Time was simply the number of min elapsed since the session began. Bars in this column show color when only the coefficient for the cumulative reward variable was significant. %Ipsi and %Contra represent the fraction of choices to ipsi- or contra-versive ports (relative to probe location in the brain) in each minute, independent of block probability. P(win-stay) is the probability of repeating the previous choice, given the previous choice was rewarded.

**Supplementary Figure 3. Histological reconstruction of recording locations.** *Left,* Histology photomicrographs for each rat (IM-657, IM-1002, IM-1003, IM-1037) from which opto-tagged dopamine cells were obtained. Red: TH-staining; green: ChR2::eYFP; blue: DAPI. Scale bars: 1mm. Numbers below each photograph indicate estimated atlas coordinates of the lowest position of the optic fiber. IM-1037 brain was sliced horizontally, so fiber track appears as a circle. *Right,* coronal atlas sections with estimated dopamine cell locations in VTA-l marked as small horizontal bars.

**Supplementary Figure 4. Identification of light-responsive units. a.** Average waveforms of optogenetically-identified dopamine neurons. Average light-evoked waveforms are shown in blue and session-wide average waveforms are in black. All spikes within 10ms of laser onset were used to construct light-evoked waveform average. **b.** Session-wide average waveform for non-dopamine cells. **c.** Opto-tagging p-value for all units plotted in log-scale, showing a strong bimodal distribution. To classify units as light-responsive we used a threshold of p<0.001. **d.** Times to first spike after laser onset, showing mean for each identified dopamine neuron, and standard deviation (jitter).

**Supplementary Figure 5. Properties of each individual identified dopamine cell** *(one per page).* **a.** Average light-evoked spike waveform (blue) and session-wide average waveform (black). **b.** Interspike interval histogram (during bandit task). **c.** Raster plot showing response to 5ms laser pulses (delivered at 2Hz). **d.** Raster plot with 10ms laser pulses (for cells that were tested under this condition). **e.** Scatter plot (as Fig. 2b), with this neuron highlighted in yellow. **f.** Behavior, and **g.** activity during the Pavlovian approach task. **h.** Firing rate, latency and reward rate during the bandit task. **i.** Average response of this cell to the bandit task *Side-In* event, broken down by reward rate terciles. **j.** Spike rasters and firing rate histograms aligned to various bandit task events.

**Supplementary Figure 6. Overall rate of VTA-l dopamine cell burst firing is not affected by reward rate. a**, Example of burst detection algorithm in action. We used a “80/160 template” approach that has long been the standard method for detecting dopamine cell bursts^57^. Each time an inter-spike-interval of 80 ms or less occurs, these and subsequent spikes are considered part of a burst until there is an interval of 160 ms or more. Numbers indicate the number of spikes in each detected burst. **b**, No change in overall rate of bursts between higher- and lower-reward rate blocks in each session. Wilcoxon paired test z=0.82, p>0.4. **c**, No change in burst rates across a wide distribution of spikes/burst. Kolmogorov-Smirnov statistic = 0.165, p>0.63). **d**, Latencies indicate substantial shifts in motivation within the same sessions. Wilcoxon paired test z=-4.28, p<1.8×10^-5^.

**Supplementary Figure 7. Distinct activity patterns of VTA-l dopamine and non-dopamine neurons.** Format is as Fig. 4, except showing both non-dopamine neurons (top) and dopamine neurons (bottom). Rasters again show one representative neuron of each class, and peri-event histograms show average for all neurons of that class. Note that the non-dopamine cells show activity during movements, starting just before *Center-In* (irrespective of latency), just before *Side-In*, just before *Food-Port-In*. For the *Light-On* versus *Center-In* comparison (scatter plot), 2-way ANOVA with factors of Latency and Alignment, Alignment F=48.9, p<0.0001, Latency n.s. F=0.82 p=0.44.

**Supplementary Figure 8. Different methods for calculating reward expectation produce similar results. a**, As Fig. 4f,g except that reward expectation was estimated using either the number of rewards in the last 10 trials (top), an actor-critic model (middle), or a Q-learning model (bottom). The two models were both trial-based, rather than evolving continuously in time. The actor-critic model estimated the overall probability of receiving a reward on each trial, V, using the update rule V’ = V + alpha (RPE), where RPE = actual reward [1 or 0] – V. The Q-learning model kept separate estimates of the probabilities of receiving rewards for left and right choices (QL, QR) and updated Q for the chosen action (only) using Q’ = Q + alpha (RPE), where RPE = actual reward [1 or 0] – Q. The learning parameter alpha was determined for each session by best fit to latencies, for V or (QL + QR) respectively. **b**, Correlations between RPE and firing were similar regardless of which method was used to estimate reward expectation.

